# Elucidating and Overcoming Lignin-Mediated Growth Suppression in Anaerobic Fungi

**DOI:** 10.64898/2026.11.26.769390

**Authors:** Shiyan Jin, Isabella R. Farrand, Wenni Zhang, Elaine Kirschke, Michelle A. O’Malley

## Abstract

Native to the gastrointestinal tract of herbivores, anaerobic gut fungi (AGF) (*Neocallimastigomycetes*) have evolved highly efficient enzymatic machinery for lignocellulose degradation. While AGF are capable of degrading lignocellulose, they more readily degrade its cellulose and hemicellulose components. Lignin is the most recalcitrant component of lignocellulose that serves as both structural support and a microbial defense strategy within plants. In this study, a negative correlation between AGF growth and substrate lignin content was quantified, ranging from pure cellulose to the lignin-rich substrate eucalyptus. Based on this observation, we investigated the impact of lignin on AGF viability and concluded that lignin delayed and slowed AGF’s growth trajectory, with IC_50_ ranging from 10.7 – 16.3 mg/mL, and specifically that lignin-rich substrates decreased overall growth. Meanwhile, lignin acted as an AGF deterrent, likely by preventing access to essential sugar-based polymers and lowering overall nutrient bioavailability. However, AGF growth was restored following glucose supplementation on otherwise growth-limiting woody substrates. Through experimental, sequential serial batch culture challenge, we successfully adapted one strain of AGF (*Neocallimastix californiae*) to grow on lignin-rich eucalyptus. This study supports the industrial valorization of lignocellulose using AGF by elucidating the relationship between the major bottleneck, lignin, on fungal growth. It also provides a strategy for optimizing fungal strains selected for growth in nutrient-deficient and lignin-rich environments.

**Importance:** Anaerobic gut fungi (AGF) are among the most powerful lignocellulose-degrading microorganisms known and represent promising workhorse for the degradation of plant biomass. However, despite their remarkable capacity to deconstruct plant cell walls, the factors limiting AGF growth on lignin-rich feedstocks remain poorly understood. This study demonstrated that lignin constrains AGF growth through both direct inhibitory effects and reduced nutrient accessibility, establishing quantitative relationships between lignin concentration and fungal viability. Furthermore, experimental evolution resulted in one AGF strain, *Neocallimastix californiae*, with improved growth on lignin-rich substrates. These findings aim to advance the understanding of lignin-mediated growth suppression and develop strategies to overcome its inhibitory effects in AGF, supporting the industrial valorization of lignocellulose.

**Research Area:** Molecular and Cellular Biology

**Key Points:**

- Anaerobic fungi preferentially degrade and uptake cellulose and hemicellulose from plant biomass compared to lignin
- Lignin has cytostatic and cytotoxicity effects on growth of anaerobic fungi
- Lignin reduces nutrient accessibility for AGF
- Experimental evolution of *Neocallimastix californiae* enables tolerance in nutrient limiting, lignin-rich environments

## INTRODUCTION

Lignocellulose, the key component of plant cell walls, is a promising feedstock for industrial chemical production due to its high availability and carbon-rich composition[1–3]. It is composed of cellulose, hemicellulose, and lignin, with cellulose being the most abundant natural biopolymers on earth[4–7]. Cellulose, the most abundant component of lignocellulose, is made of D-glucose monomers linked by β(1→4)-bonds[1]. Additionally, hemicellulose, a branched polysaccharide, has a composition varies by plant type, with hardwoods rich in xylan and softwoods rich in glucomannan[8]. A number of enzymes have been identified and engineered that efficiently degrade both cellulose and hemicellulose, leading to more effective conversion of lignocellulosic biomass into biofuels, biochemicals, and other value-added products.

While carbohydrate-based cellulose and hemicellulose have established industrial conversion pathways, lignin stands out as a heterogeneous aromatic polymer with untapped potential for producing high-value aromatic chemicals[6]. Lignin is formed from three subunits: minor p-hydroxyphenyl (H), predominant guaiacyl (G), and syringyl (S) that is particularly enriched in hardwood[9]. It is a major natural source for manufacturing aromatic compounds. However, the various bonding patterns of lignin across plant types present major barriers for efficient depolymerization and downstream valorization into commodity chemicals [9–11]. The low-molecular-weight breakdown products generated from lignin depolymerization comprise a diverse array of aromatic compounds, such as vanillin, syringaldehyde and other phenolic derivatives[9]. Nevertheless, the heterogeneous composition of these product mixtures may pose challenges for downstream processing and industrial utilization.

Despite their potential as the next-generation biofuels and chemical feedstocks, lignocellulosic materials are notoriously difficult to valorize due to their complex structure[6,10–12]. Efficient deconstruction typically requires multiple catalytic steps under harsh physical conditions, rendering the process both environmentally and economically expensive[13,14]. Microbial bioprocessing offers an attractive alternative, as many microbes have evolved to deconstruct biomass under benign conditions[15]. For instance, anaerobic gut fungi (AGF), classified under the phylum *Neocallimastigomycota*, play a key role in the digestive tract of herbivores[15–18]. Their unique growth environment has driven the evolution of specialized traits, such as rhizoid morphologies for substrate penetration and enzyme complexes that localize carbohydrate-active enzymes (CAZymes) for efficient lignocellulose deconstruction[17,19,20].

While AGF are highly effective at breaking down polysaccharide cellulose and hemicellulose, lignin presents a significant obstacle[5]. As a complex aromatic polymer, lignin functions as a defense barrier against pathogens and is thereby difficult to biologically deconstruct[5,9,21]. Known microbial lignin-degrading enzymes, such as laccases and peroxidases, are primarily found in aerobic fungi and require oxygen to generate radical intermediates that are critical for lignin deconstruction[22]. These oxygen-dependent mechanisms are presumed to be inoperative in anaerobic environments. Nevertheless, herbivore gut communities have been shown to displace lignin from biomass to access fermentable carbohydrates, indicating that some anaerobic organisms have evolved alternative mechanisms for lignin transformation[5,23,24]. Recent studies using two-dimensional heteronuclear single quantum coherence nuclear magnetic resonance (2D-HSQC-NMR) spectrometry have provided the first direct evidence of native lignin bond disruption by anaerobic fungi[5], challenging the long-standing assumption that lignin breakdown is an exclusively aerobic process.

Although AGF genomes encode a rich arsenal of CAZymes for effective and rapid biomass breakdown[17,25–27], they lack recognizable homologs of canonical lignin-degrading enzymes[5]. This is likely due, in part, to the large fraction of poorly annotated putative genes within *Neocallimastigomycota* genomes, a consequence of high A-T content, dense genomic repeats, and phylogenetic divergence from other fungal species[16]. Combined with the oxygen dependence of known lignin-degrading enzymes[6], this suggests that AGF may employ previously unrecognized, oxygen-independent mechanisms to alter the natural lignin structure.

While herbaceous crops are efficiently utilized by AGF owing to their anaerobic lignin bond cleavage and diverse CAZyme production, they are challenged by the recalcitrant nutrient barrier and antimicrobial properties of lignin [9,34–36]. In contrast to grass materials, the elevated lignin content and highly branched lignin architecture of woody substrates, such as poplar, result in poor AGF growth when they are used as the sole carbon source[5]. Experimental evolution is a powerful technique that enhances microbial performance by mimicking natural selection process to derive desired phenotypes in the laboratory[37,38]. In experimental evolution, microorganisms are subjected to progressively stringent selection pressures, promoting beneficial mutations across the population[39]. Even without an elucidated understanding of the underlying genotype-phenotype relationship, experimental evolution improves organisms, allowing the investigation of diverse stress tolerances and growth optimization in non-standard environments[39–44].

In this study, we quantified how differences in lignin concentration across representative AGF substrates impact its proliferation and growth, and elucidated two challenges associated with lignin hinderance on fungal growth. We investigated whether the detrimental effects of lignin on AGF arise from direct cytotoxicity and/or indirect nutrient limitation. To overcome lignin-mediated growth suppression, we applied experimental evolution to obtain a strain of *Neocallimastix californiae* that better tolerates eucalyptus, a hardwood biomass with a high lignin content, and subsequently compared the apparent growth of the wild-type *N. californiae* and evolved strains. Overall, this work aims to advance the understanding of lignin-mediated growth suppression and develop strategies to overcome its inhibitory effects in AGF.

## MATERIALS AND METHODS

### Routine cultivation of Neocallimastix californiae

Anaerobic Medium C (MC)[18] was used for both routine culture and growth experiments. Anaerobic fungal isolate, *N. californiae*[45], was cultivated in 10 mL MC under anaerobic conditions at 39°C in Hungate tubes with a 100% CO_2_ headspace. The substrate used for routine culture maintenance was 1% (w/v) 2 mm milled reed canary grass (RCG) (*Phalaris arundinacea*). To maintain viable and active cultures, fungal strains were routinely passaged using a syringe and needle every 3-7 days by transferring 1 mL of an actively growing culture into 9 ml of fresh MC containing RCG in anaerobic Hungate tubes.

### Growth characterization of *N. californiae* across a range of substrates

To assess fungal growth on various carbon sources, cultures were supplemented with 1% (w/v) of the following substrates: Avicel (Sigma-Aldrich, catalogue number 11363), cellobiose (Fisher Scientific, catalogue number A1455322), Whatman filter paper (Cytiva Whatman, Part No. 1001-325), sorghum (*Sorghum bicolor*), RCG, poplar (*Populus*), pine (*Pinus*), and eucalyptus (*Eucalyptus*). Whatman paper was cut into approximately 5*10 mm strips. All lignocellulosic biomass substrates provided by the Joint BioEnergy Institute, and were milled to a particle size of 2 mm using a Thomas-Wiley Mini Mill (Model 3383-L10, A. H. Thomas). Following inoculation, the cultures were maintained in 10 mL of anaerobic MC medium in Hungate tubes.

*N. californiae* develop biofilms through rhizoid structures, rendering traditional optical density (OD) measurements unsuitable for assessing growth[25]. Therefore, to evaluate fungal growth, the increase in pressure within an anaerobic Hungate tube, resulting from the production of hydrogen and carbon dioxide during metabolism, is monitored as an indicator of fungal growth[25,29]. Fermentation gas accumulation was recorded using a pressure transducer connected to a syringe and needle, and the headspace pressure of the sample was vented to 1.0 psig after each measurement to prevent pressure buildup in the Hungates. Each condition was grown in biological triplicates, alongside one blank media bottle.

The growth curve was fitted using the Gompertz model, a mathematical equation commonly employed to describe the growth of various germinating microbes, including *Aspergillus niger* and anaerobic microbial community in digestors[31,32]. This model is characteristic for its asymmetry, capturing an initial slow growth phase due to spore formation and hyphal development, followed by a rapid increase during mycelial expansion, and a subsequent deceleration as the population approaches carrying capacity[33]. The model was simplified in this study to emphasize the exponential growth phase through the growth rate coefficient (r), the inflection time (t_i_) when the fastest growth occurs, as well as the asymptote (P_max_) (Equation 1).

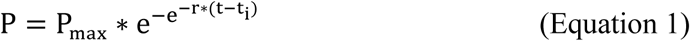

### Quantifying the effect of alkaline lignin and nutrient availability on anaerobic fungal growth

Initial aliquots of 9 mL MC medium were dispensed into Hungate tubes and supplemented with 10 mg/mL cellobiose or RCG and various concentrations of dealkaline lignin (0 mg/mL, 2 mg/mL, 5 mg/mL, 10 mg/mL, 15 mg/mL, 20 mg/mL, 25 mg/mL) prior to autoclaving. Dealkaline lignin was purchased from Fisher Scientific (TCI America, Part No. L0045). Upon inoculation, 1 mL of an actively growing *N. californiae* culture from a single 80 mL batch culture growing on either cellobiose or RCG was added to each Hungate tube to reach a total volume of 10 mL. Fermentation pressure was monitored daily over the growth period, and the tubes were vented to 1.0 psig after each measurement, as described above. Following the addition of lignin but prior to AGF inoculation, Hungate tubes containing MC medium were vented four times over a week to release accumulated gas and maintain stable culture conditions. The bicarbonate buffer system within the medium responded to pH shifts by releasing CO_2_, and venting was performed to prevent subsequent pressure increases prior to inoculation. Each condition was grown in biological triplicates, alongside one blank media bottle with respective lignin amount to account for abiotic pressure changes.

The cytotoxicity effects of lignin were modeled using the half-maximal inhibitory concentration (IC_50_) model, a parameter widely used to quantitatively describe the potency of an inhibitory compound. IC_50_ values are commonly estimated by fitting dose–response data to sigmoidal inhibition models and provide a convenient framework for comparing inhibitor sensitivities across biological systems[46,47]. The fitted upper asymptote (P_max_) represents the maximum growth attainable in the presence of lignin, while the lower asymptote (P_min_) represents the minimum growth predicted by the model. The Hill slope (H_hill_) characterizes the sensitivity of fungal growth to changes in lignin concentration (C_lignin_) around the IC₅₀ value (Equation 2). In this study, IC_50_ was defined as the lignin concentration required to reduce fungal growth by 50% relative to the fitted upper asymptote (P_max_) of the sigmoidal dose-response curve.

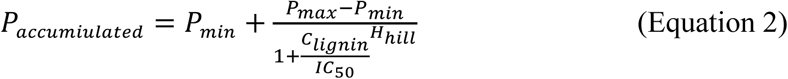

To further characterize the impact of nutrient availability on fungal growth, initial aliquots of 9 mL of MC medium were dispensed into Hungate tubes and supplemented with 10 mg/mL of eucalyptus or pine prior to autoclaving. Upon inoculation, 1 mL of an actively growing *N. californiae* culture from a single 80 mL batch culture growing on RCG was added to each Hungate tube using a syringe to reach a total volume of 10 mL. Fermentation pressure was monitored daily over a 12-day period using a pressure transducer, and the tubes were vented to 1.0 psig after each measurement, as described above. On Day 4, 1 mL of 10 mg/mL sterile glucose solution or sterile water was added to all Hungates. A single uninoculated control containing corresponding substrate was included to account for abiotic pressure changes.

### Microscopy and characterization of morphological changes in *N. californiae*

Initial aliquots of 9 mL MC medium were dispensed into Hungate tubes containing 10 mg/mL cellobiose. For lignin-treated cultures, kraft lignin was supplemented to a final concentration of 2 mg/mL prior to autoclaving. *N. californiae* cultures were inoculated into the Hungate tubes for 3 days under anaerobic conditions. Fungal cells were fixed, and their morphology was examined using an inverted light microscope.

### Serial batch cultivation of *N. californiae* to establish lignin tolerance

Experimental evolution was performed to improve fungal growth on lignin-rich biomass. For experimental evolution, 9 mL of MC medium was prepared and aliquoted into Hungate tubes with 100 mg of total biomass composed of a mixture of reed canary grass and eucalyptus in varying proportions. The percentage of eucalyptus ranged from 0% to 100% in 10% increments per stage. At each composition stage, the cultures were incubated anaerobically for 3-4 days before passage. Three serial transfers were conducted at each eucalyptus proportion by inoculating 1 mL of an actively growing culture into fresh medium. Evolved *N. californiae* strains were maintained on 100% eucalyptus substrates for five passages before subsequent characterization experiments and cryopreservation. Two *N. californiae* strains were evolved in parallel, then cryopreserved by mixing cultures with sterile glycerol to a final concentration of 11.5% (v/v) in MC medium. Samples were stored at −80 °C for long-term preservation.

Following the completion of experimental evolution, the evolved *N. californiae* strain was compared with the wild-type *N. californiae* for growth performance on 10 mg/mL of each RCG, eucalyptus, and pine. For each substrate, 9 mL MC medium was inoculated with 1 mL actively growing culture in Hungate tubes. The fermentation pressure was recorded and vented to 1.0 psig daily for six days, as described above.

### Statistical analysis

In this study, statistical analyses were performed on biological triplicates. Statistical comparisons were done using two-tail Student’s T-test, unless noted otherwise. The error bars included in the study represent the standard deviations of three biological triplicates, unless otherwise mentioned.

## RESULTS AND DISCUSSION

### *N. californiae* is proficient at decomposing diverse lignocellulosic biomass

*N. californiae*, a filamentous AGF strain originally isolated from the feces of goats, has adapted to metabolize lignocellulose derived from crude biomass. The strain is capable of utilizing various types of substrates, ranging from soluble sugars to complex lignocellulose. While generating ATP as cellular energy, they produce hydrogen and carbon dioxide during the same processes, resulting in increased pressure[25,28–30]. Fungal growth was well modeled using the Gompertz model (Equation 1) with R-square values above 0.99 for most substrates (Fig. 1A, Table S1), where the fitted parameters are shown in Table S1.

**Figure 1.**
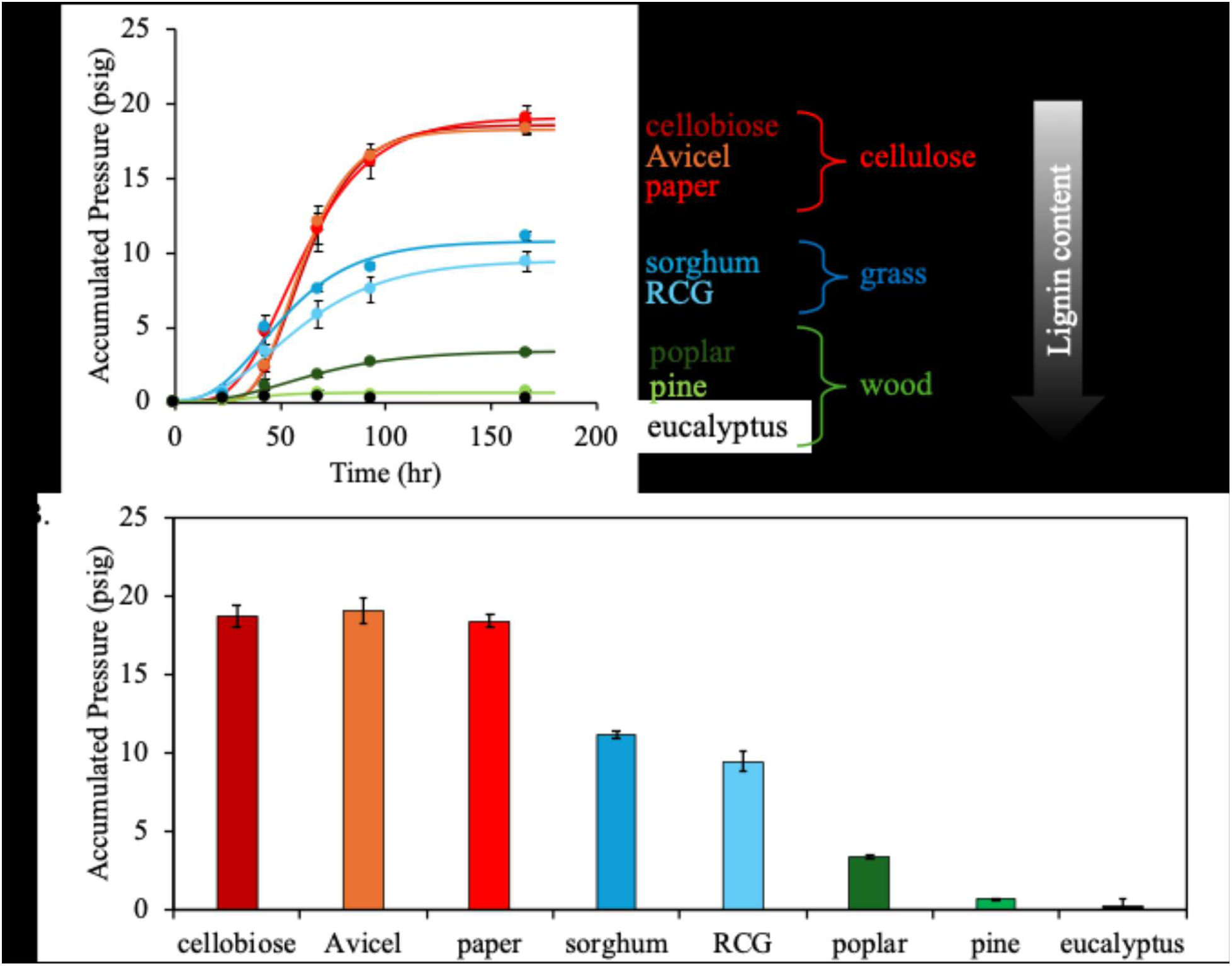
*N. Californiae* growth exhibits a negative correlation against increasing lignin content in representative substrates. **A.** Time-course of pressure accumulated over 7 days when *N. californiae* grown on different substrates. Higher growth was observed on cellulose, followed by grass and then wood, indicating that increasing lignin content is associated with reduced fungal growth. Growth was fitted using the Gompertz model (Eqn. 1). **B.** Endpoint pressure accumulation corresponding to the final time point (Day 7) from the same experiment shown in A. Substrate tested were cellobiose, Avicel, paper, sorghum, reed canary grass (RCG), poplar, pine and eucalyptus. Error bars represent standard deviations of biological triplicates (n = 3).

*N. californiae* reached the highest final pressure, indicating the best growth on cellulose substrates, including soluble cellobiose, crystalline Avicel, and fibrous filter paper, with final pressure of 18.7 ± 0.7 psig, 19.1 ± 0.8 psig, 18.4 ± 0.4 psig, respectively. The morphological form of cellulose did not significantly influence the final growth titers of *N. californiae*, as all three types achieved consistent total pressure over a 7-day period (Fig. 1A, 1B).

In addition to utilizing simple sugars such as cellulose, *N. californiae* can grow on un-pretreated biomass as its sole carbon source. However, their growth on grass materials, such as sorghum and RCG was less robust compared to cellulose, as indicated by the lower overall pressure accumulated (Fig. 1A, 1B). Additionally, fungal growth was notably impeded when exposed to more woody, lignin-rich substrates such as poplar, pine, and eucalyptus (Fig. 1A, 1B). The variation in growth performance is likely attributed to the presence of lignin, which impedes fungal development and proliferation. Compared to grass, woody lignocellulose is characterized by a higher lignin content, which likely results in a higher dosage of toxic metabolized lignin byproducts, such as vanillic acid[48]. Furthermore, lignin layers encasing the cellulose core may limit sugar accessibility for fungi, leading to overall detrimental growth on woody substrates that generally possess a higher lignin content. Overall, these results indicate that lignin hinders the growth of AGF, even though these fungi have developed the ability to decompose raw plant material. This is consistent with the role of lignin in protecting plants from microbial invasion. In this study, we proposed and assessed two possible mechanisms by which lignin might restrict fungal development: either by releasing toxins into the medium or by obstructing access to the underlying nutrients for fungal proliferation.

### Lignin supplementation induces growth suppression and influences rhizoid development in *N. californiae* growing on cellobiose

To investigate the impact of lignin on fungal growth, dealkaline lignin, despite having fewer β-O-4 linkages[53,54], was introduced as substitute for natural lignin into an anaerobic fungal culture growing on cellobiose, a simple soluble sugar. The addition of lignin at concentrations below 2 mg/mL did not affect the final pressure accumulated, suggesting that fungi can tolerate low lignin concentrations without compromising significant growth (Fig 2A, 2B). Conversely, when lignin concentration exceeded 5 mg/mL, a negative impact on fungal growth was observed, as suggested by a reduction in the accumulated pressure. Furthermore, a pronounced cytotoxic effect was noted when more than 15 mg/mL of lignin was added, resulting in no recorded accumulated pressure within AGF cultures. (Fig. 2A, 2B).

**Figure 2.**
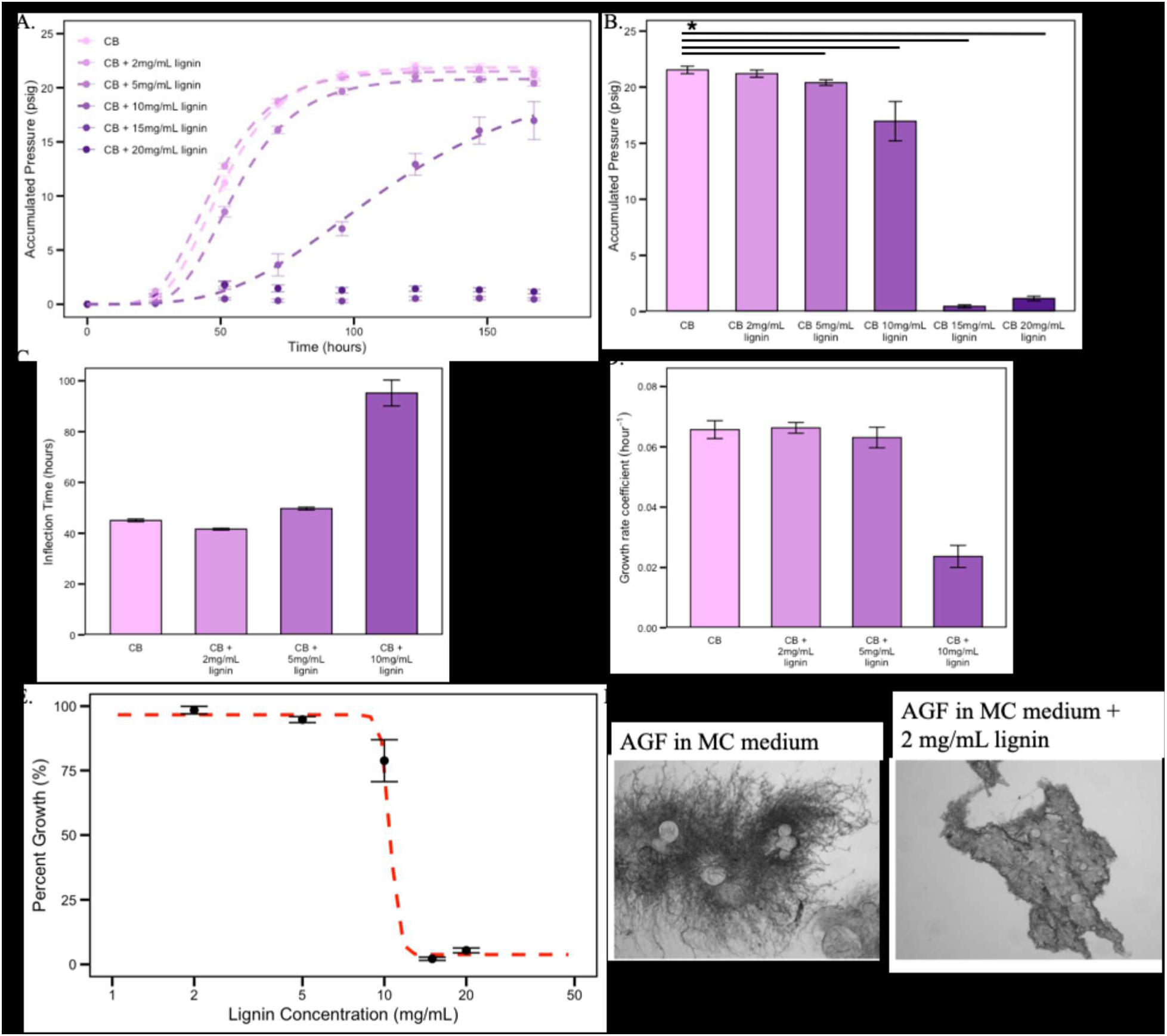
Lignin supplementation hinders *N. californiae* growth on cellobiose. **A.** Time-course of pressure accumulation by *N. californiae* grown on cellobiose (CB) with 0 mg/mL, 2 mg/mL, 5 mg/mL, 10 mg/mL 15 mg/mL, and 20 mg/mL of lignin added. Fungal growth was fitted using the Gompertz model (Eqn. 1). **B.** Endpoint pressure accumulated by *N. californiae* after 7 days of growth under the conditions shown in panel A. Statistical significance between groups is indicated by * and connecting horizontal lines (p < 0.1, two-tailed Student’s t-test). **C.** Infection time parameters derived from Gompertz model fitting at different lignin concentrations. **D**. Growth rate coefficients derived from Gompertz model fitting at different lignin concentrations. **E.** Dose–response analysis using an IC50 model (Eqn. 2) to quantify the effect of lignin on fungal growth on CB and estimate lignin-induced growth inhibition. The fitted parameters shown in Table S4. F. – G. Morphological changes under microscope of *N. californiae* grown on CB in **(F.)** MC medium alone or **(G.)** MC medium supplemented with 2 mg/mL lignin. For panels A, B, and E, error bars represent standard deviations from biological triplicates (n = 3). For panels C and D, error bars represent standard deviations from model-derived parameter estimates. Cellobiose is abbreviated as CB.

Supplementing lignin not only reduced the overall pressure buildup associated with AGF growth, but it also altered the growth trajectory. The addition of dealkaline lignin resulted in a general downward and rightward shift in the growth curve (Fig. 2A). Introducing less than 5 mg/mL of dealkaline lignin did not significantly alter the overall growth curve, as modeled by the Gompertz equation, which returned similar inflection times (t_i_) and growth rate coefficients (r) for samples with 2 mg/mL, and 5 mg/mL lignin (Fig. 2C, 2D). However, the addition of more than 10 mg/mL of lignin led to a pronounced shift of the growth curve, supported by a significant higher t_i_, smaller r, and lower P_max_ compared to the control without any added lignin (Fig. 2C, 2D. Table S2). This delay suggests that fungi initiate the exponential growth phase later, corroborating the cytostatic effect of dilute lignin.

When the lignin concentration exceeded 10 mg/mL, no fungal growth was observed, suggesting that the concentration reached the cytotoxicity threshold. To quantitatively assess the cytotoxicity effect, the IC_50_ model was employed, which determined that10.7 mg/mL of lignin as the concentration to achieve 50% of maximum growth with the presence of lignin (Fig. 2E, Table. A4). The steep Hill slope suggests that lignin induced multimodal cytotoxicity, as multiple distinct mechanisms or pathways were impacted simultaneously. This is further supported by the induced morphological changes in *N. californiae* (Fig. 2F, 2G). Sensing the pretense of lignin triggers response likely reprograms fungal growth behavior under lignin-rich conditions and promoting cell clustering.

We hypothesized that the majority of the change in growth performance observed here arises from the cellular response from lignin addition, rather than the introduced carbon from dealkaline lignin, since no cell growth was observed on lignin alone, indicating that lignin cannot serve as the sole carbon source to sustain a *N. californiae* culture under these culture conditions (Fig. S1). Moreover, the MC medium utilized in the study possesses sufficient buffering capacity to maintain a biologically suitable pH when lignin was added at concentrations below 25 mg/mL, thereby preventing potential cell death caused by pH fluctuations (Fig. S2).

### Enhanced lignin tolerance in *N. californiae* grown on RCG

The observation that lignin induces morphological changes in fungi suggests that it initiates specific cellular responses, potentially promoting the decomposition of more complex substrates. In comparison to soluble cellobiose, fungi have been observed to secrete a more diverse array of proteins for the lignocellulose degradation, such as CAZymes[17,20], suggesting that lignin’s presence alone might trigger a complex cellular reaction beyond its possible toxic effects. Therefore, the interaction between the added dealkaline lignin and secreted lignocellulolytic proteins was investigated using insoluble lignocellulose RCG as a comparison to soluble cellobiose, with the former requiring a more diverse enzyme composition to achieve hydrolysis. The previously observed negative correlation between lignin and cell growth was conserved on RCG substrates, as evidenced by the decrease in overall pressure accumulated with increased lignin added (Fig. 3A, 3B). Nevertheless, AGF was more sensitive to lignin when growing on RCG, as a smaller dose of 2 mg/mL of lignin was sufficient to trigger a significant reduction in fungal growth, leading to a 51.4% decrease in pressure accumulation, opposite of the lack of difference in growth observed for 2 mg/mL lignin on cellobiose. This implies that the variation in fungal growth could also be attributed to the inhibition of protein by lignin and its breakdown products. This sequestration effect could lead to an initial inhibitory impact when the lignin concentration is low[55].

**Figure 3.**
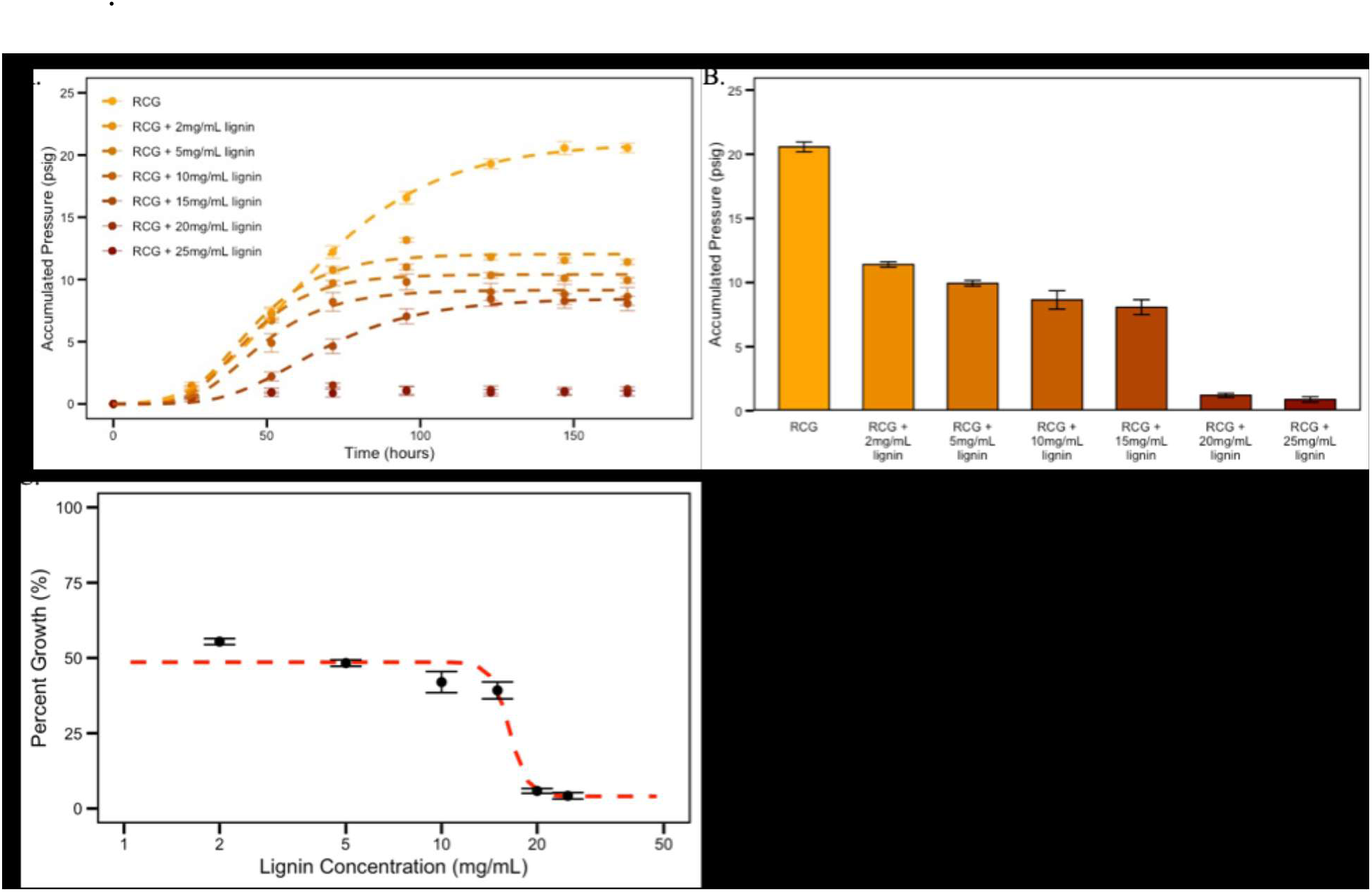
Lignin exhibits cytotoxic effects on *N. californiae* when cultivating with reed canary grass (RCG). **A.** Time-course of pressure accumulation by *N. californiae* grown on RCG with 0 mg/mL, 2 mg/L, 5 mg/mL, 10 mg/mL, 15 mg/mL, 20 mg/mL, and 25 mg/mL of lignin added. Fungal growth was fitted using the Gompertz model (Eqn. 1). **B.** Endpoint pressure accumulation by *N. californiae* after 7 days of growth under the conditions shown in panel A. **C.** Dose–response analysis using an IC50 model (Eqn. 2) to quantify the effect of lignin on fungal growth in RCG and estimate lignin sensitivity. The fitted parameters shown in Table S4. Error bars represent standard deviations from biological triplicates (n = 3). Reed canary grass is abbreviated as RCG.

Although fungi were sensitive to the additional of dealkaline lignin, the AGF cytotoxicity threshold for lignin when growing on RCG was 20 mg/mL, higher than that for cellobiose, suggesting greater lignin tolerance (Fig. 3A, 3B). Notably, two distinct growth response ranges suggest that lignin can exert a cytostatic effect at lower concentrations, inhibiting the fungal growth, while triggering profound cytotoxicity effect at higher levels. To quantify these effects, the IC_50_ was calculated to be 16.3 mg/mL of dealkaline lignin added to the RCG (Fig. 3C). Notably, both IC_50_ values, from cultures grown on cellobiose and RCG, fall in the range of other literature values of 0.4 to 27.7 mg/mL[36,56]. Additionally, the Gompertz model suggested the asymptote percent growth is 48.6% on RCG substrate, significantly lower than the 96.6% growth observed on cellobiose. It is worth mentioning that in woody biomass, such as pine, lignin constitutes approximately 20% - 25% of the total weight (Table 1) [51], equivalent to 2 mg/mL - 2.5 mg/mL of lignin in 10 mL Hungate culture tested. However, prominent fungal death was found only at higher lignin concentration, above 10 mg/mL of kraft lignin (Fig. 3A-C), suggesting that, in addition to the presence of lignin, multiple factors contribute to cell death on woody substrates.

**Table 1.** Lignin composition of substrates evaluated in this study.

| Substrate type | Substrate | Lignin Content (%) | Reference |
|---|---|---|---|
| Cellulose | cellobiose, Avicel, paper | 0 | [10] |
| Grasses | sorghum, RCG | 17 – 24 | [49,50] |
| Hardwood | poplar, eucalyptus | 20 – 25 | [49,51,52] |
| Softwood | pine | 28 – 32 | [49,52] |

### High lignin content in woody substrates restricts fungal penetration and access to underlying cellulose and hemicellulose

A reduction in AGF growth on woody substrates, such as poplar, compared to cultures grown on cellulose and grass substrates, was observed (Fig. 1A). The plausibility of the poor nutrient hypothesis is further supported by the restored fungal growth on substrates that are typically uninhabitable after supplementing extra sugars. To examine the limited growth observed on woody substrates such as eucalyptus and pine, glucose was introduced into the culture on the fourth day to provide additional sugars to promote growth. While AGF displayed no initial growth, pressure started to accumulate once glucose was introduced into the system four days after the fungi were placed on the two substrates (Fig 4). This observation indicates that the lignin concentration in the solution was not sufficiently toxic to eradicate the fungi completely. Instead, it appears to impede fungal access to the cellulose and hemicellulose core, ultimately resulting in fungal mortality from insufficient sugars.

**Figure 4.**
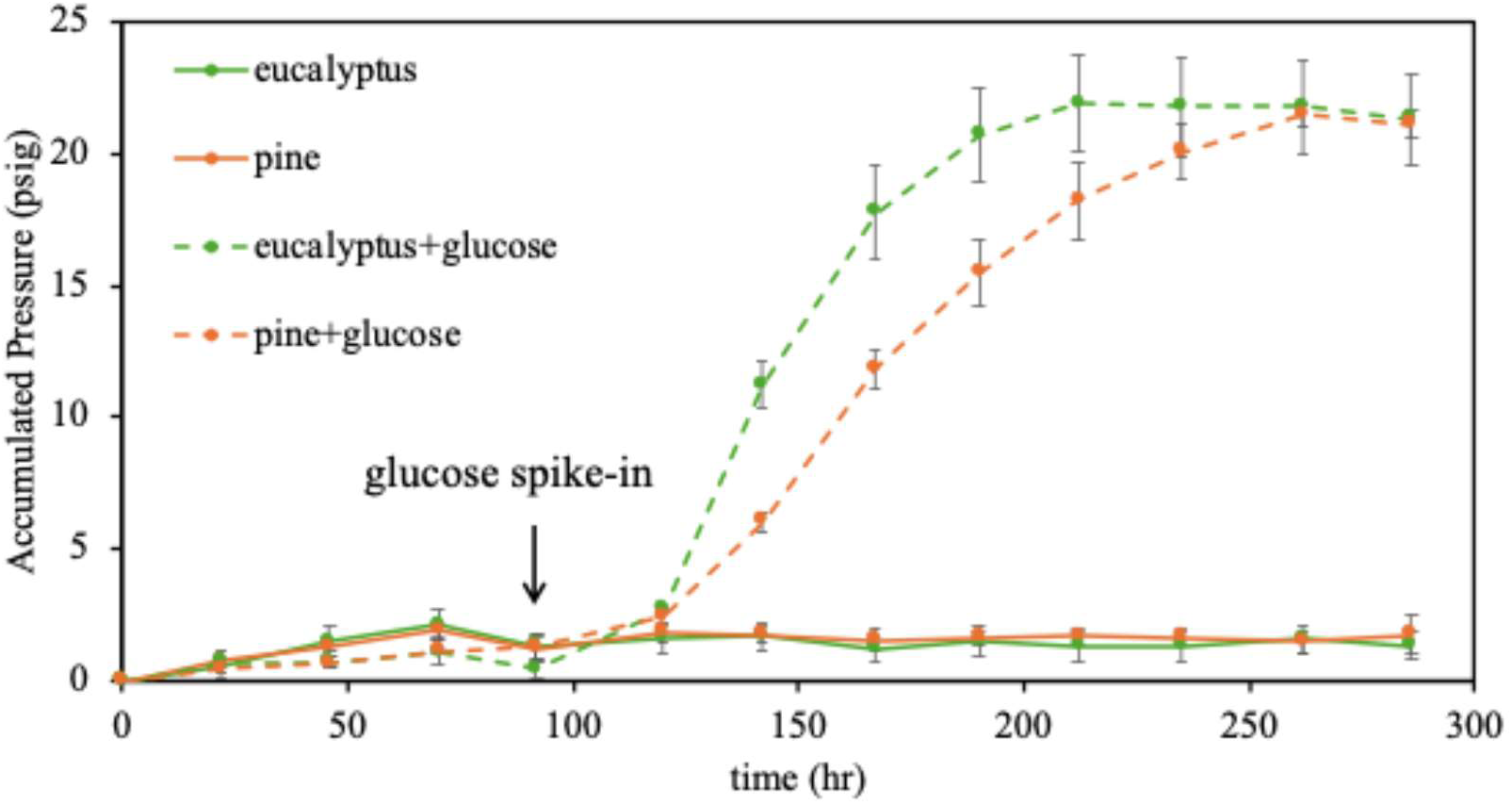
Time course of pressure accumulated by *N. californiae* when growing on traditionally inhabitable eucalyptus and pine, with or without later supplementation of 1 mg/mL glucose as an additional carbon source. Pressure was monitored over 12 days. Each condition was performed in biological triplicates (n = 3), and error bars represent the standard deviations the triplicate measurements.

### Serial batch cultivation to evolve a more robust lignin-tolerant AGF strain

An experimental evolution study was conducted to develop an *N. californiae* strain with enhanced growth capabilities on substrates with high lignin content, which typically inhibits fungal growth. The study gradually exposed *N. californiae* to increasing concentrations of eucalyptus, which is a more lignin-rich substrate compared to RCG (Fig. 5A, Table 1). The gradual decrease in nutrient availability was designed to induce mutations or epigenetic modifications in *N. californiae*, facilitating its adaptation to nutrient-deficient conditions. The results show a general decline in the growth of *N. californiae* as the concentration of eucalyptus increased (Fig. 5B).

**Figure 5.**
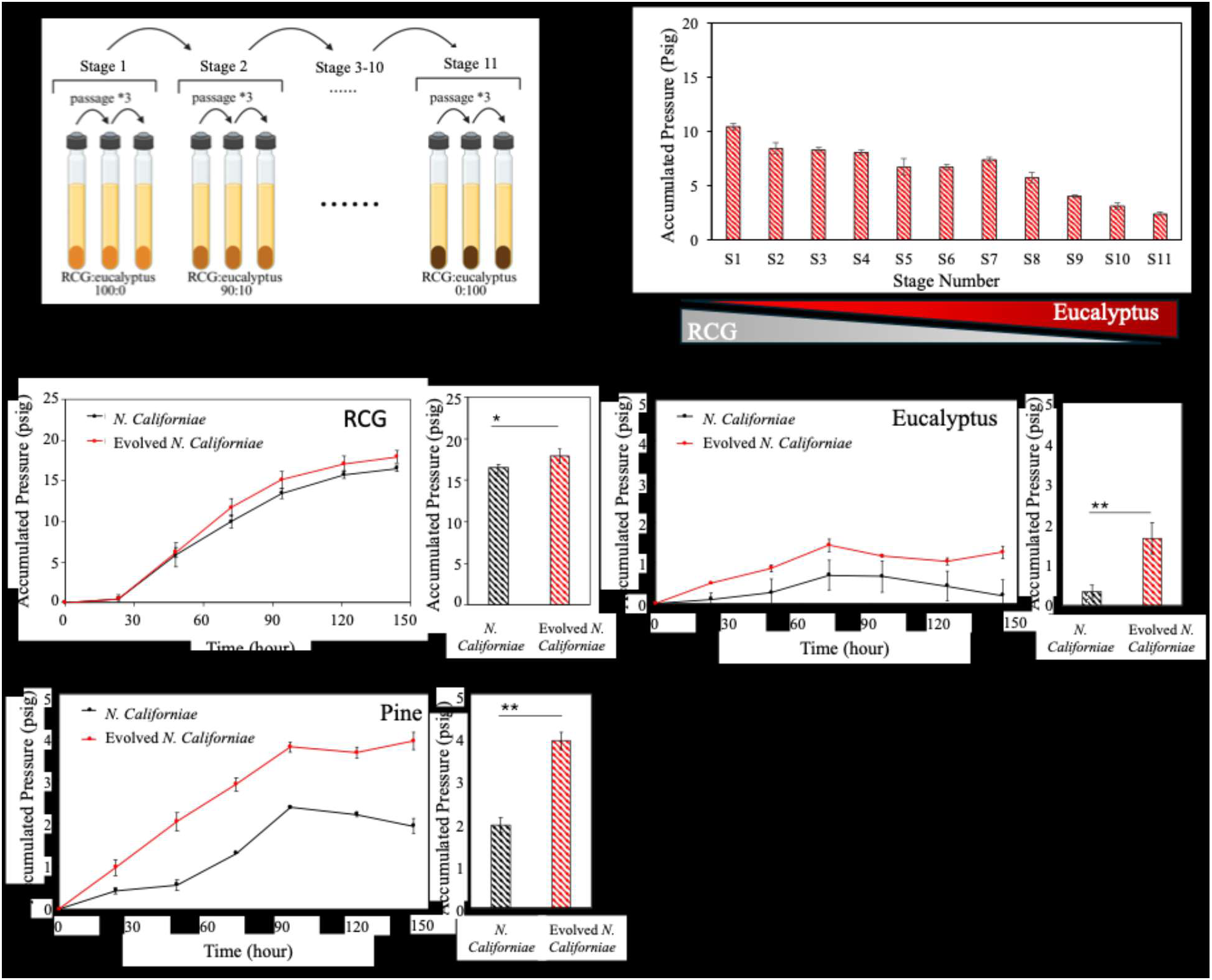
Experimental evolution of *N. californiae* yields an evolved strain with enhanced growth on plant substrates. **A.** Schematic illustration of the experimental evolution setup, where RCG:eucalyptus ratio ranged from 100:0 to 0:100. For each stage, eucalyptus ratio to RCG increased by 0.1, and over 11 stages reached 1. At each stage, three passages were made at the same RCG to eucalyptus ratio. Panel A was created using BioRender.com. **B.** Monitoring of the experimental evolution process, tracked by assessing fungal growth through pressure accumulation at each stage throughout the evolution experiment. **C.** Time-course pressure accumulation of *N. californiae* and evolved N. californiae strains grown on RCG. **D.** Endpoint pressure accumulation after 6 days on RCG, showing higher growth of the evolved strain compared to *N. californiae*. **E.** Time-course pressure accumulation of *N. californiae* and evolved strains grown on eucalyptus. **F.** Endpoint pressure accumulation after 6 days on eucalyptus, showing higher growth of the evolved strain compared to *N. californiae*. **G.** Time-course pressure accumulation of *N. californiae* and evolved strains grown on pine. **H.** Endpoint pressure accumulation after 6 days on pine, showing higher growth of the evolved strain compared to *N. californiae*. Statistical significance is indicated as *p < 0.1 and **p < 0.05. Reed canary grass is denoted as RCG. For A., error bars represent standard deviation from the two co-evolved strains across the three passages at each stage (n = 6). For C. – H., error bars represent standard deviations from biological triplicates (n = 3).

After a three-month evolution, two strains of *N. californiae* were been developed in parallel that could consistently grow using eucalyptus as their sole carbon source. To evaluate the phenotypic changes caused by experimental evolution, the evolved strains were assessed by growth in different nutrient level environments compared to the native *N. californiae* strain. When cultivated on RCG, both native and evolved *N. californiae* strains exhibited robust growth, with the evolved strain demonstrating modestly superior performance compared to that of the wild-type *N. californiae* (Fig. 5C, 5D). In contrast, when *N. californiae* was subjected to traditionally challenging substrates, such as eucalyptus and pine, the evolved strain, although still underperforming relatively to RCG, exhibited enhanced robustness compared to the native *N. californiae* after six days (Fig. 5E-5H). The accumulated pressure for evolved strain was threefold higher than that of the *N. californiae* on eucalyptus and twofold higher on pine. The higher pressure accumulated across all three substrates suggests that the evolved strain generally exhibited greater growth robustness and resilience in nutrient-poor environments compared to the wild-type *N. californiae*, indicating that mutations and/or epigenetic changes in the strain occurred as a consequence of the experimental evolution.

Unlike model microorganisms such as *E. coli*, whose genomes have been extensively annotated and for which mutations arising during experimental evolution can be readily identified, anaerobic gut fungi AGF possess highly complex genomes that remain poorly characterized. Approximately 70% of the genome of *N. californiae* remains unannotated, reflecting substantial uncertainty regarding gene content and function[17]. In addition, AGF genomes are highly A-T rich and contain extensive repetitive regions, creating significant challenges for high-quality DNA extraction, genome assembly, and accurate variant calling[16]. As a result, mutations arising during experimental evolution would likely occur within poorly annotated regions, and distinguishing true mutations from sequencing or assembly artifacts would be difficult. Therefore, no genomic analysis was performed as part of this experimental evolution study.

## CONCLUSION

In this study, we demonstrated that although AGF have the advantage of growing on crude lignocellulosic biomass, the amount of lignin present can adversely affect fungal growth. Two challenges induced by lignin are either lignin is lethal to the cells, or it significantly impedes AGF accessibility to the underlying sugars. We tested these challenges and concluded that the lignin, as well as associated toxic hydrolysates, can impede fungal growth, with the extent depending on the complexity of substrates used. Additionally, the lignin layer encasing the biomass can present a formidable barrier to fungal penetration. Furthermore, we conducted an experimental evolution study to evolve *N. californiae* to survive in nutrient-scarce environments. An evolved strain was successfully developed that utilized eucalyptus as the only carbon source and exhibited a more robust growth phenotype than that of the *N. californiae*. This experimental evolution framework provides a valuable pipeline for enhancing lignocellulosic substrate utilization in AGF and related fungal systems. Future studies focusing on the specific genetic mutations of the evolved strain relative to the wild type would be beneficial in elucidating the genes responsible for detoxification, growth, and the production of key proteins essential for growth.

## STATEMENTS AND DECLARATIONS

### Data Availability Statement

All relevant data generated and analyzed in this study are included in this article or the supplementary information.

## Acknowledgements

We are grateful to the researchers at JBEI for providing ground biomass substrates for the study. Figure 5A was generated using BioRender.com.

## Funding

This material is based upon work supported by the Joint BioEnergy Institute, U.S. Department of Energy, Office of Science, Biological and Environmental Research Program under Award Number DE-AC02-05CH11231. The work conducted at the Joint BioEnergy Institute was supported by the U.S. Department of Energy, Office of Science, Biological and Environmental Research Program, through contract DE-AC02-05CH11231 between Lawrence Berkeley National Laboratory and the U.S. Department of Energy.

### Author Contributions

M.A.O, and S.J. conceived the idea and designed the experiments. S.J. carried out most of the experiments, analyzed the data, and participated in the manuscript writing. W. Z. helped collecting data for the lignin response study. I.R.F. helped collecting data for the experimental evolution study. E.K. helped taking microscopic images. M.A.O. supervised the project.

### Ethical approval

This article does not contain any studies with human or animal participants performed by any of the authors.

### Supplemental Materials

**Table S1.**
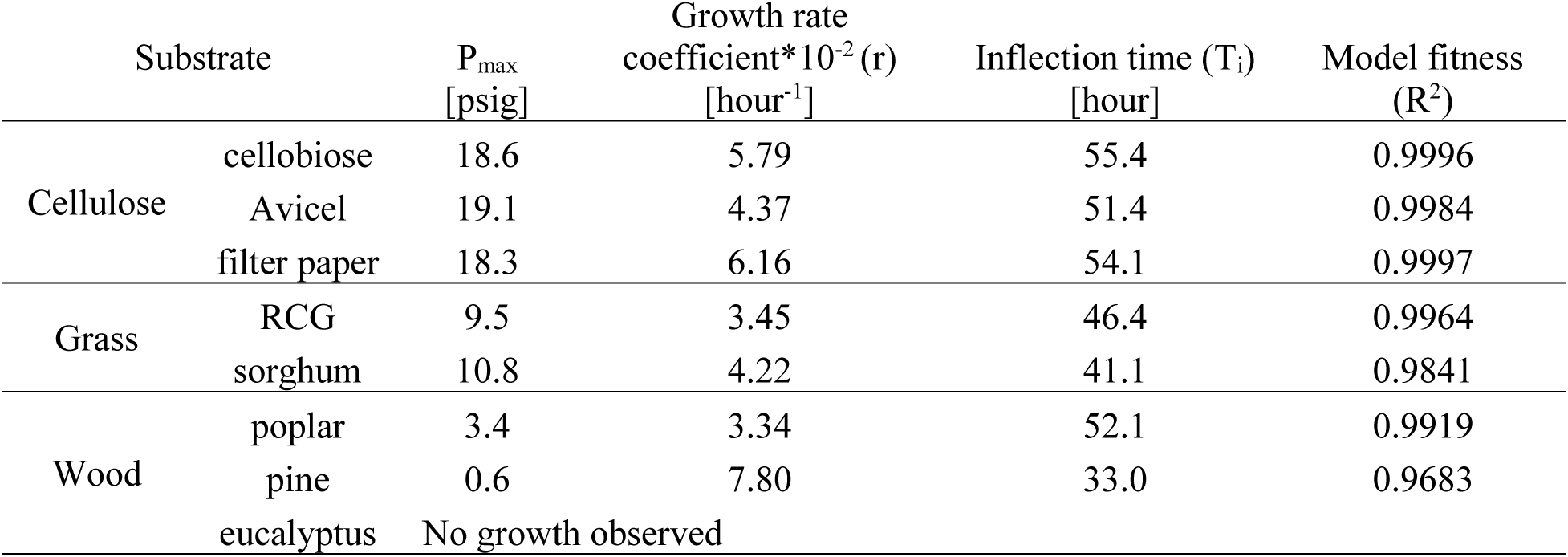
Gompertz model parameters for Figure 1

| Substrate |  | P<sub>max</sub><br>[psig] | Growth rate coefficient*10<sup>-2</sup> (r)<br>[hour<sup>-1</sup>] | Inflection time (T<sub>i</sub>)<br>[hour] | Model fitness<br>(R<sup>2</sup>) |
|---|---|---|---|---|---|
| Cellulose | cellobiose | 18.6 | 5.79 | 55.4 | 0.9996 |
|  | Avicel | 19.1 | 4.37 | 51.4 | 0.9984 |
|  | filter paper | 18.3 | 6.16 | 54.1 | 0.9997 |
| Grass | RCG | 9.5 | 3.45 | 46.4 | 0.9964 |
|  | sorghum | 10.8 | 4.22 | 41.1 | 0.9841 |
| Wood | poplar | 3.4 | 3.34 | 52.1 | 0.9919 |
|  | pine | 0.6 | 7.80 | 33.0 | 0.9683 |
|  | eucalyptus | No growth observed |  |  |  |

**Table S2.** Gompertz model parameters for Figure 2A

| Substrate | P<sub>max</sub><br>[psig] | Growth rate coefficient*10<sup>-2</sup><br>(r) [hour<sup>-1</sup>] | Inflection time<br>(T<sub>i</sub>) [hour] | Model fitness<br>(R<sup>2</sup>) |
|---|---|---|---|---|
| CB | 21.88 | 6.57 | 45.04 | 0.9995 |
| CB + 2 mg/mL lignin | 21.51 | 6.63 | 41.63 | 0.9998 |
| CB + 5 mg/mL lignin | 20.80 | 6.31 | 49.71 | 0.9994 |
| CB + 10 mg/mL lignin | 20.88 | 2.37 | 95.18 | 0.9953 |

**Table S3.** Gompertz model parameters for Figure 3A.

| Substrate | P<sub>max</sub> [psig] | Growth rate<br>coefficient*10<sup>-2</sup> (r) [hour<sup>-1</sup>] | Inflection time<br>(T<sub>i</sub>) [hour] | Model fitness<br>(R<sup>2</sup>) |
|---|---|---|---|---|
| RCG | 21.02 | 3.61 | 54.17 | 0.9991 |
| RCG + 2 mg/mL lignin | 12.04 | 6.36 | 39.04 | 0.9824 |
| RCG + 5 mg/mL lignin | 10.41 | 7.08 | 38.54 | 0.9919 |
| RCG + 10 mg/mL lignin | 9.15 | 6.40 | 41.70 | 0.9829 |
| RCG + 15 mg/mL lignin | 8.48 | 4.37 | 58.41 | 0.9941 |

**Table S4.** IC50 model parameters for Figure 2E and 3C

| Parameters | Figure 2E | Figure 3C |
|---|---|---|
| P<sub>max</sub> | 96.6% | 48.6% |
| P<sub>min</sub> | 3.4% | 4% |
| Hill slope | 28.4 | 15.3 |
| IC<sub>50</sub> | 10.5 mg/mL | 16.3 mg/mL |

**Figure S1.**
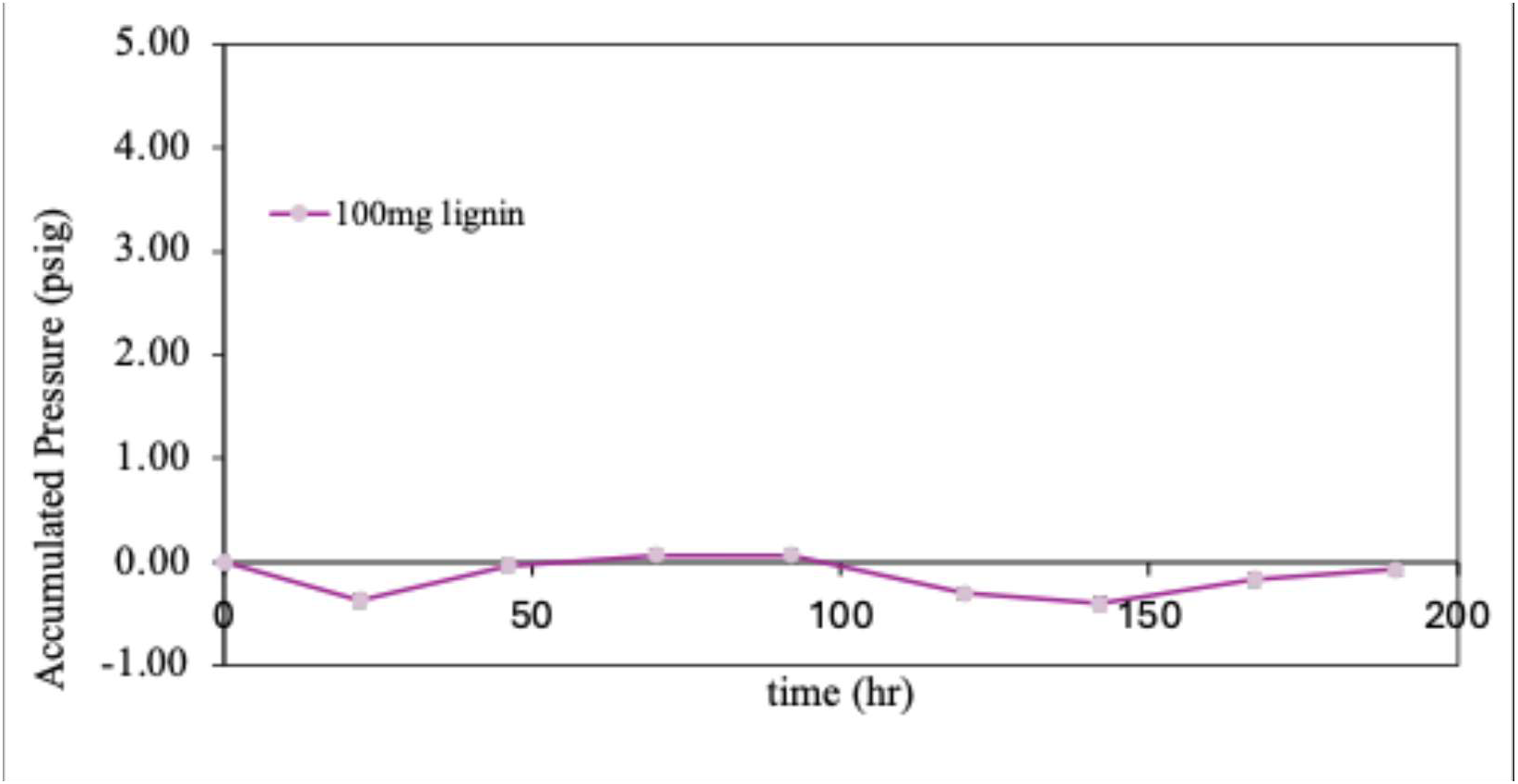
Time-course of accumulated pressure when growing *N. californiae* on lignin

**Figure S2.**
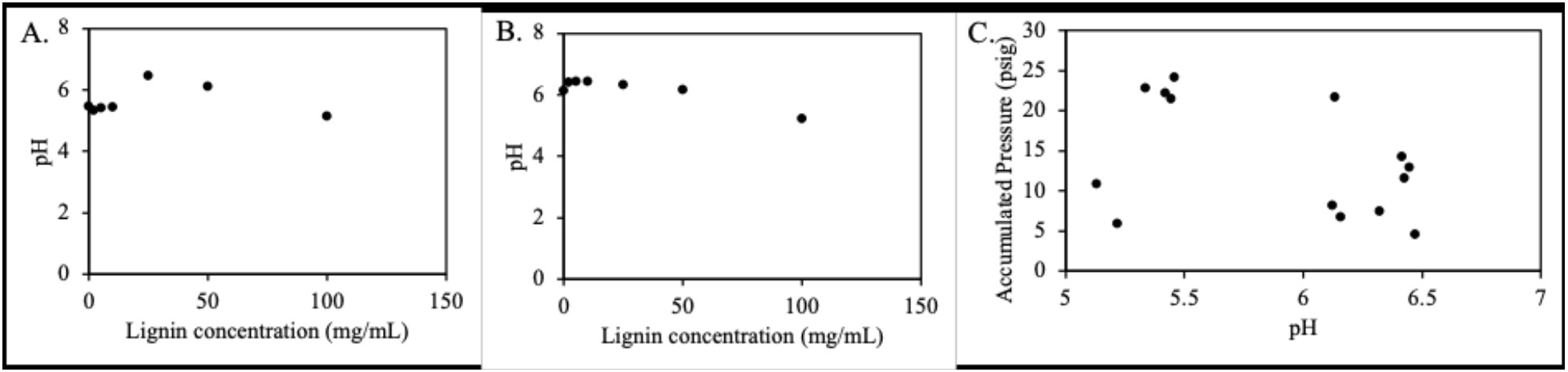
Lignin did not change the medium pH enough to change the growth pattern

